# Nitric oxide synthase-mediated early nitric oxide-burst alleviates drought-induced oxidative damage in ammonium supplied-rice roots

**DOI:** 10.1101/383323

**Authors:** Cao Xiaochuang, Zhu Chunquan, Zhong Chu, Zhang Junhua, Zhu Lianfeng, Wu Lianghuan, Ma Qingxu, Jin Qianyu

## Abstract

Ammonium (NH_4_^+^) can enhance rice drought tolerance in comparison to nitrate (NO_3_^-^). The mechanism underpinning this relationship was investigated based on the time-dependent nitric oxide (NO) production and its protective role in oxidative stress of NH_4_^+^-/NO_3_^-^-supplied rice under drought. An early burst of NO was induced by drought 3h after root NH_4_^+^ treatment but not after NO_3_^-^ treatment. Root oxidative damage induced by drought was significantly higher in NO_3_^-^ than in NH_4_^+^-treatment due to its reactive oxygen species accumulation. Inducing NO production by applying NO donor 3h after NO_3_^-^ treatment alleviated the oxidative damage, while inhibiting the early NO burst increased root oxidative damage in NH_4_^+^ treatment. Application of nitric oxide synthase (NOS) inhibitor N(G)-nitro-*L*-arginine methyl ester (L-NAME) completely suppressed NO synthesis in roots 3h after NH_4_^+^ treatment and aggravated drought-induced oxidative damage, indicating the aggravation of oxidative damage might have resulted from changes in NOS-mediated early NO burst. Drought also increased root antioxidant enzymes activities, which were further induced by NO donor but repressed by NO scavenger and NOS inhibitor in NH_4_^+^-treated roots. Thus, the NOS-mediated early NO burst plays an important role in alleviating oxidative damage induced by drought by enhancing antioxidant defenses in NH_4_^+^-supplied rice roots.

**Highlight:** NOS-mediated early NO burst plays an important role in alleviating oxidative damage induced by water stress, by enhancing the antioxidant defenses in roots supplemented with NH_4_^+^

---

**Abbreviations**
(APX): ascorbate peroxidase
(CAT): catalase
(c-PTIO): 2-(4-carboxyphenyl)-4,4,5,5-tetramethylimidazoline-1-oxyl-3-oxide
(L-NAME): N(G)-nitro-*L*-arginine methyl ester
(MDA): malondialdehyde
(NO): nitric oxide
(NOS): nitric oxide synthase
(NR): nitrate reductase
(ONOO^-^): peroxynitrite
(PEG-6000): polyethylene glycol
(POD): peroxidase
(RNS): reactive nitrogen species
(ROS): reactive oxygen species
(SNP): sodium nitroprusside
(SOD): superoxide dismutase

## Introduction

As human population and global climate change increase, drought stress is becoming a major abiotic factor limiting crop growth and yield. Plants have evolved several strategies to contend with water stress. These include morphological,physiological, and molecular adaptations (Bogeat-Triboulot *et al.,* 2007; Guo *et al.,* 2007; Slewinski, 2012). Nitric oxide (NO) is an important signaling molecule in various physiological functions like seed germination, floral transition, stomatal movement, leaf senescence, and yield development, and it has gained increasing attention since the 1980s (Neill *et al.,* 2003; Wilson *et al.,* 2007; Simontacchi *et al.,* 2015). Certain plant responses and adaptations to abiotic stresses involve NO, and sufficient data indicate that NO mediates plant responses to various stimuli including drought (Mata and Lamattina, 2001), salt (Zhao *et al.,* 2007), and metal toxicity (Gonzalez *et al.,* 2012) stresses, thereby enhancing plant stress tolerance and survival.

Water deficits significantly increase NO production in plants (Signorelli *et al.,* 2013; Planchet *et al.,* 2014). As a free radical, NO can form various reactive nitrogen species (RNS) such as peroxynitrite (ONOO^-^), nitrogen dioxide (NO2), dinitrogentrioxide (N2O3) and S-nitrosoglutathione (GSNO), which are involved in many physiological functions of plants (del Rio, 2015), indicating that NO and NO-derived molecules take part in inorganic nitrogen (N) metabolism. A combination of transgenic technology and pharmacological analysis have indicated that NO induces antioxidant activity and alleviates water stress in plants in several ways: 1) It limits reactive oxygen species (ROS) accumulation and ROS-induced cytotoxic activity by inhibiting the ROS-producer NADPH oxidase via S-nitrosylation (Fan *et al.,* 2012); 2) It functions as an antioxidant and reacts with ROS (e.g. O_2_^-^) to generate transient ONOO^-^, which is then scavenged by other cellular processes (del Rio, 2015); 3) It induces the expression of genes coding for antioxidant enzymes, such as superoxide dismutase (SOD), ascorbate peroxidase (APX), and glutathione reductase (GR), and may increase enzyme activity by posttranslational modifications thereby reducing lipid peroxidation under water stress (Farooq *et al.,* 2009; Fan and Liu, 2012); 4) It helps maintaining high vacuolar concentrations of osmotically active solutes and amino acids like proline (Verdoy *et al.,* 2006); and 5) It acts as a downstream abscisic acid (ABA) signal molecule and participates in “ABA-H_2_O_2_-NO-MAPK” signal transduction processes. It also increases plant antioxidant ability (Zhang *et al.,* 2007). The accumulation of ROS in water-stressed plants impairs the function of biochemical processes, damages organelles, and ultimately results in cell death (Jiang and Zhang, 2002). Therefore, endogenous NO production may enhance plant antioxidant capacity and help plant cells survive under various types of stresses.

However, NO also has biphasic properties on plants. The duality of its effects depends on stress duration and severity, and on the cell, tissue, and plants species (Neill, 2007; Santisree *et al.,* 2015). At low concentration or early stage of abiotic stress, NO participates in important functions in higher plants through its involvement in physiological and stress-related processes (as described above). Arasimowicz-Jelonek *et al*. (2009a, b) demonstrated that NO synthesis slightly increased in roots subjected to <10 h water deficit, but significantly up-regulated after prolonged (>17h) drought. Under severe or protracted longtime stress, NO overproduction in plants can shift the cellular stress status from oxidative stress to severe nitrification stress, finally damaging proteins, nucleic acids, and membranes (Groß *et al.,* 2013; del Rio 2015). Protein tyrosine nitration is considered a good marker to evaluate the process of nitrosative stress under various abiotic environments (Corpas *et al.,* 2007, 2008). Excess NO can also act synergistically with ROS resulting in nitro-oxidative stress and eliciting undesirable toxic effects in plant cells (Signorelli *et al.,* 2013). Liao *et al*. (2012) and Sun *et al*. (2014) argued that the ability of endogenous or exogenous NO production in plants to alleviate oxidant damage was dose-dependent. Therefore, determining instantaneous plant NO content under drought stress may not completely reflect the specific role of NO in drought tolerance.

In higher plants, nitrate reductase (NR) and nitric oxide synthase (NOS) are the two key enzymes for NO production (Guo *et al.,* 2003; Neill *et al.,* 2003). Moreover, NR-dependent NO production occurs in response to pathogen infection (Shi and Li, 2008), drought (Freschi *et al.,* 2010), and freezing (Zhao *et al.,* 2009). Arasimowicz-Jelonek *et al*. (2009a, b) applied the NO donor sodium nitroprusside (SNP) and GSNO to water-stressed cucumbers and demonstrated that both NR and NOS participated in drought tolerance. Shi *et al*. (2014) reported that rat neuronal NO synthase overexpression in rice plants increased their tolerance to drought stress, thus demonstrating the importance of NOS-mediated NO production in water deficit tolerance. Despite increasing knowledge on NO-mediated plant functions, NO origins and signaling in response to prolonged stress and their regulation in plant drought tolerance remain poorly understood.

Ammonium (NH_4_^+^) and nitrate (NO_3_^-^) are the two primary N sources for plants. It is known that the negative effects of drought stress on plant development can be more effectively alleviated by NH_4_^+^ than NO_3_^-^ supplementation, as evaluated by plant growth, physiological characteristics, and gene expression levels (Guo *et al.,* 2007,Yang *et al.,* 2012; Ding *et al.,* 2015). NO has a key role in plant water stress acclimation and drought tolerance. Nevertheless, information on the dynamic changes in NO production and its role in drought acclimation in plants supplied with NO_3_^-^ or NH_4_^+^ during the early stages of water stress is scarce. In the present study, variations in endogenous NO production were monitored in roots supplied with this two N nutrition supplements during water stress. The specific role and origin of the endogenous NO produced were investigated using pharmacological methods. The present study revealed that an early NO burst is crucial for alleviating the water stress-induced oxidative damage through enhancement of antioxidant defenses in roots of NH_4_^+^-supplied plants. Further analyses demonstrated that this early NO burst might be triggered by NOS-like enzymes.

## Materials and methods

### Plant material and growth conditions

Rice (*Oryza sativa* L. ‘Zhongzheyou No. 1’ *hybrid indica*) seedlings were grown hydroponically in a greenhouse. Seeds were sterilized in 1% (v/v) sodium hypochlorite solution. After germination, seeds were transferred to a 0.5 mmol L^-1^ CaCl_2_ solution (pH 5.5). Three days later, the seedlings were transferred to 1.5-L black plastic pots containing a solution with the following composition: NH_4_NO_3_ (0.5 mM), NaH_2_PO_4_·,2H_2_O (0.18 mM), KCl (0.18 mM), CaCl_2_(0.36 mM), MgSO4·7H_2_O (0.6 mM), MnCl2·4H_2_O (9 μM), Na2MoO4·4H_2_O (0.1 μM), H3BO3 (10 μM), ZnSO4·7H_2_O (0.7 μM), CuSO4 (0.3 μM), and FeSO4·7H_2_O-EDTA (20 μM). All experiments were performed in a controlled growth room under the following conditions: 14/10 h light/dark photoperiod, 400 μmol m^-2^ s^-1^ light intensity, 28°C or 23°C during day or night, respectively, and 60% relative humidity. The solution pH was adjusted to 5.5 with 5 mM2-(N-Morpholino)ethanesulfonic acid (MES). The solution was replaced every 3 days.

After 6 days, seedlings of similar size were cultivated under one of the following treatments: 1 mM NO_3_^-^, 1 mM NO_3_^-^ + 10% polyethylene glycol (PEG-6000), 1 mM NH_4_^+^, or 1 mM NH_4_^+^ + 10% PEG-6000. Water stress was induced by adding 10% PEG-6000. Eight treatments were performed in the NO donor (i.e., SNP) experiments: NH_4_^+^, NH_4_^+^+ SNP, NH_4_^+^+ PEG-6000, NH++ PEG-6000 + SNP, NO_3_^-^, NO_3_^-^+ SNP, NO_3_^-^+ PEG-6000, and NO_3_^-^+ PEG-6000 + SNP. The final SNP concentration was 20 M. For each N nutrition experiment, treatments receiving sufficient water were defined as control (CK) treatments.

For the NO scavenger2-(4-carboxyphenyl)-4,4,5,5-tetramethylimidazoline-1-oxyl-3-oxide (c-PTIO, 100 μM) experiment, rice seedlings supplied with 1 mM NO_3_^-^ or 1 mM NH_4_^+^ solution were pretreated with c-PTIO for 3 h and then given sufficient water (CK) or subjected to water stress for 24 h under the same conditions as those described above.

To investigate the effects of the NO biosynthesis inhibitors, rice seedlings supplied with 1 mM NO_3_^-^ or 1 mM NH_4_^+^ solution were pretreated with the NO scavenger, tungstate NR inhibitor (100 μM), or NOS inhibitor [Nx-Nitro-*L*-arginine methyl ester hydrochloride (L-NAME); 100 μM] for 3 h, and then given sufficient water (CK) or subjected to water stress for 24 h under the same conditions as described above. There were eight treatments for each N nutrition: Tungstate, L-NAME, Tungstate + SNP, PEG-6000 + Tungstate, PEG-6000 + Tungstate + SNP, L-NAME + SNP, PEG-6000 + L-NAME, and PEG-6000 + L-NAME + SNP.

### Determination of NO and ONOO^-^contents

The 4-amino-5-methylamino-2’,7’-difluorofluorescein diacetate (DAF-FM DA) probe was used to determine endogenous root NO levels (Sun *et al.,* 2014). Root tips (1 cm) were incubated with 10 μM DAF-FM DA in the dark for 30 min, washed 3× with 20 mM HEPES-KOH (pH 7.4) to remove excess fluorescence, and then observed and photographed under a Nikon Eclipse 80i fluorescence microscope (Nikon, Tokyo, Japan; EX 460-500, DM 505, BA 510-560). The relative fluorescence intensity was measured with Photoshop v. 7.0 (Adobe Systems, Mountain View, CA, USA).

Root endogenous ONOO^-^ was determined using the aminophenylfluorescein (APF) probe method. Root tips were incubated with 10 μM APF dissolved in 10 mM Tris-HCl (pH 7.4) in the dark for 60 min, and then washed 3× with 10 mM Tris-HCl. Fluorescence images and relative fluorescence intensities were analyzed as described above for NO.

### Histochemical analyses

Lipid peroxidation and root cell death were histochemically detected with Schiff’s reagent and Evans blue (Yamamoto *et al.,* 2001). Root tips were incubated in Schiff’s reagent for 20 min and washed by three consecutive immersions in 0.5% (w/v) K_2_O_3_S solution. A red/purple endpoint indicated the presence of aldehydes generated by lipid peroxidation. Roots were also washed by performing three serial immersions in distilled water, then incubated in 0.25% (w/v) Evans blue for 15 min, and finally washed 3× with distilled water. Roots stained with Schiff’s reagent and Evans blue were immediately photographed under a Leica S6E stereomicroscope (Leica, Solms, Germany).

The oxidative damage level, specifically expressed as membrane lipid peroxidation and protein oxidative damage, were estimated by measuring the concentrations of malondialdehyde (MDA) and carbonyl group with 2,4-dinitrophenylhydrazine (DNPH) according to the methods described in Velikova *et al*. (2000).

### Determination of ROS contents

Root O_2_^-^ content was estimated using the method described in Liu *et al*. (2007) with some modifications: about 0.15 g fresh root was powdered with 2 mL of 65 mM phosphate buffer saline (PBS, pH 7.8) in a pre-cooled mortar, and centrifuged at 5,000 × *g* for 10 min at 4 °C. Then, 0.9 mL of 65 mM PBS (pH 7.8) and 0.1 mL of 10 mM hydroxylammonium chloride were added to 1 mL of the root extract supernatant, thoroughly mixed, and left to react for 25 min. After this period, 1 mL of 1% (w/v) sulfanilamide and 1 mL of 0.02% (w/v) *N*-(1-naphthyl)-ethylenediaminedihydrochloride were added to 1 mL of root extract solution and left to react for 30 min. Absorbance was then measured at 540 nm.

Root H_2_O_2_ content was determined by the photocolorimetric method: ~0.15 g fresh root was powdered with 2 mL acetone in a pre-cooled mortar, and centrifuged at 5,000 × *g* for 10 min at 4 °C. Then, 0.1 mL of 5% (w/v) TiSO_4_ and 0.1 mL pre-cooled ammonium hydroxide were added to 1 mL of the root extract supernatant, which was re-centrifuged at 5,000 × *g* for 10 min. The supernatant was discarded and the sediment was re-dissolved in 4 mL of 2 M H2SO4. The absorbance of the root extract solution was measured at 415 nm (Wang *et al.,* 2010).

Root OH^-^ was analyzed by the methods described in Liu *et al*. (2010): ~0.1 g fresh root was powdered with 3 mL of 50 mM PBS (pH 7.0) in a mortar, and centrifuged at 10,000 × *g* for 10 min at 4 °C. Then, 1.0 mL of 25 mM PBS (pH 7.0) containing 5 mM 2-deoxy-.D-ribose and 0.2 mM NADH were added to 1 mL of the root extract supernatant, completely blended, and left to react for 60 min at 35°C in the dark. Following this incubation, 1 mL of 1% (w/v) thiobarbituric acid and 1 mL glacial acetic acid were added to the filtrate. The mixture was heated to 100°C for 30 min and then placed on ice for 20 min. The absorbance of the root extract solution was then measured at 532 nm, and the OH^-^ content was inferred from the production of MDA.

### Determination of enzyme activities

Fresh rice root samples (0.5 g) were homogenized in 5 mL of 10 mM phosphate buffer (pH 7.0) containing 4% (w/v) polyvinylpyrrolidone and 1 mM ethylenediaminetetraacetic acid. The supernatant was used as crude enzyme solution and collected by centrifugation at 12,000 × *g* for 15 min at 4°C. The activities of SOD, catalase (CAT), APX, and peroxidase (POD) were estimated using the photocolorimetric methods described in Jiang and Zhang (2002) and Sachadyn-Krol *et al*. (2016).

Root NR and NOS activities were assayed using the methods described in Scheible *et al*. (1997) and Lin *et al*. (2012), with some modifications. Briefly, total protein was extracted using a buffer containing 100 mM HEPES-KOH (pH 7.5), 1 mM EDTA, 10% (v/v) glycerol, 5 mM DTT, 0.5 mM phenylmethylsulfonyl fluoride, 0.1% Triton X-100 (v/v), 1% PVP, and 20 μM FAD. The supernatant was collected by centrifugation at 12,000 × *g* for 20 min at 4°C, and then used to determine the NR and NOS activities at 520 nm and 340 nm, respectively.

### Determination of arginine and citrulline

Arginine and citrulline contents were estimated using the method described in Salazar *et al*. (2012). Briefly, 1.0 g root samples were frozen in liquid N2 and extracted with 4 mL 80% (v/v) methanol, and then centrifuged at 10,000 × *g* for 5 min at 4 °C. The supernatant was then used in derivatization and reaction processes. Serial concentrations of amino acid standards were prepared as described above for the derivatizing reagent, and the derivatizing samples were used to determine the arginine and citrulline contents using liquid chromatography/electrospray ionization tandem mass spectroscopy(LC-ESI-MS).

### Statistical analyses

All experiments conducted in this study were performed in triplicate, at least. All data, expressed as means ± standard error (SE), were processed in SPSS v. 13.0 (IBM Corp., Armonk, NY, USA). The Least Significant Difference (LSD) test was used to determine statistical significant differences among the treatments (P<0.05). Figures were drawn in Origin v. 8.0 (OriginLab Corporation, Northampton, MA, USA).

## Results

### Plant growth and physiological characteristics

Growth-and physiology-related parameters, such as biomass, photosynthesis rate (*P_n_*), and root N uptake rate in rice seedlings supplied with different N sources were negatively and differently influenced by the 21days water stress (Supplementary Fig. S1a-f). While there were significant decreases in the biomass of NO_3_^-^-supplied plants (62.1% and 52.2% reductions in shoot and total biomass, respectively) (Supplementary Fig. S1a, c), biomass accumulation was not significantly affected in NH_4_^+^-supplied plants, in relation to CK plants. Water stress reduced *Pn* in the leaves of NO_3_^-^-treated plants by 40.4% (P<0.05) but that of NH_4_+-treated plants was only reduced by 17.3% (Supplementary Fig. S1d) in relation to CK plants. Thus, NH_4_^+^-supplied rice seedlings can alleviate PEG-induced drought stress more effectively than NO_3_^-^-supplied rice seedlings.

### Root endogenous NO production and histochemical analyses of oxidative damage

To investigate whether NO participates in water stress acclimation, endogenous NO levels in the roots were monitored with the NO-specific fluorescent probe DAF-FM DA. Significant differences in endogenous NO production were observed in roots after 48 h of water stress (Fig. 1a). In CK plants, NO production was relatively stable and varied little between the two N treatments. In contrast, water stress significantly induced endogenous NO production 3 h after the roots received NH_4_^+^. However, endogenous NO gradually increased only after 6 h in the NO_3_^-^ treatment. Relative fluorescence indicated a significant early burst of NO at 3 h of water stress in the NH_4_^+^ treatment relative to the control. The NO level in the seedlings treated with NH_4_^+^ was 2.92× higher than that ofNO_3_^-^-treated plants. Nevertheless, NO in the NO_3_^-^-treated seedlings was 2.72×higher than in NH_4_^+^-treated plants after 24 h of water stress (Fig. 1b).

**Fig. 1.**
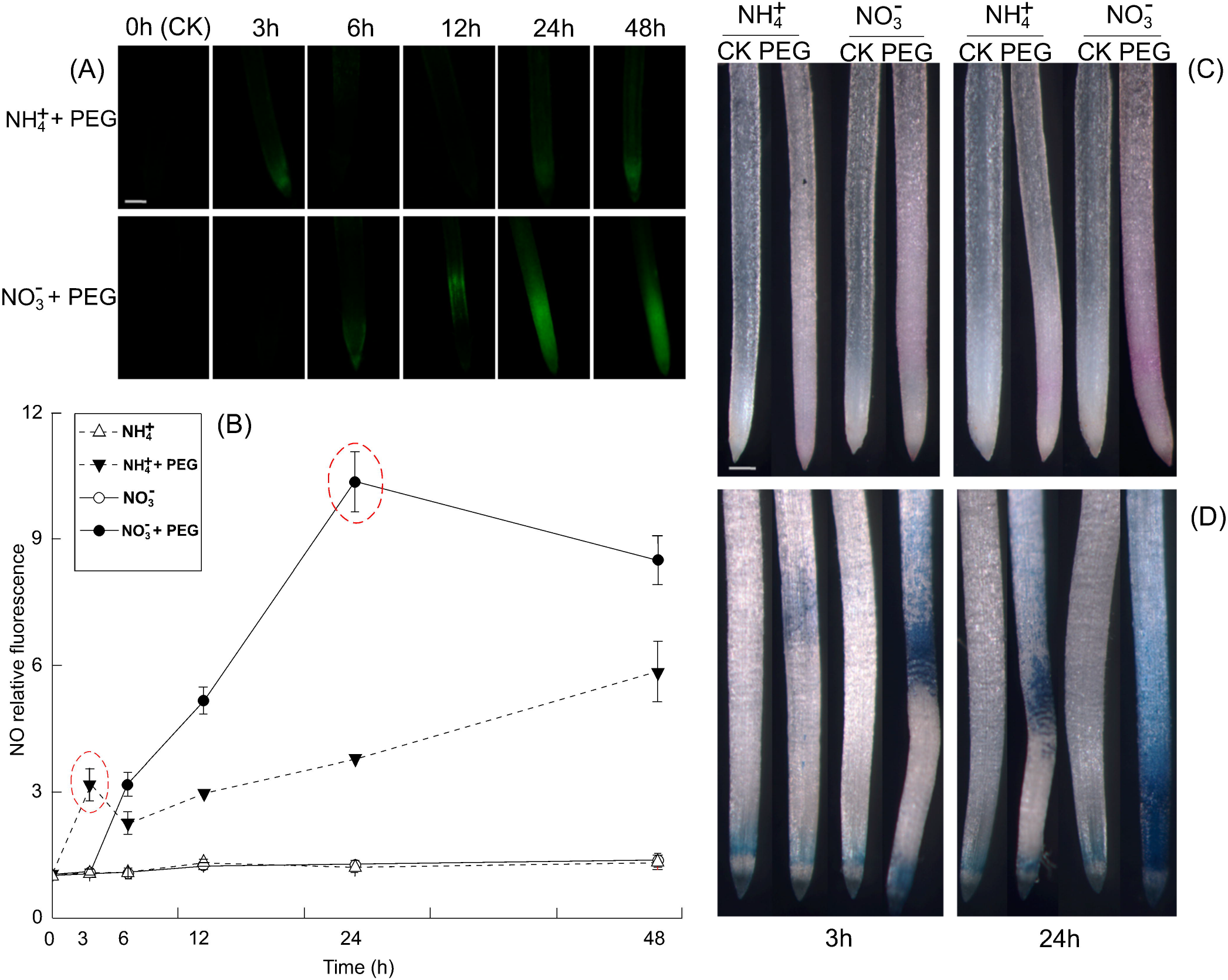
Time-dependent endogenous nitric oxide (NO) production and histochemical detection of oxidative damage in the root apices ofNH_4_^+^- and NO_3_^-^-supplied rice seedlings under water stress. (a) Detection of NO fluorescence using 4-amino-5-methylamino-2’,7’-difluorofluorescein diacetate (DAF-FM DA) staining and a fluorescence microscope. NO generation is indicated by green fluorescence. Bar=300 μm. (b) NO production is expressed as relative fluorescence. To detect the NO production time course, seedling roots exposed to 10% polyethylene glycol(PEG) were collected at 0, 3, 6, 12, 24, and 48 h. (c) and (d) Histochemical detection of the aldehydes derived from lipid peroxidation and Evans blue uptake in root apices of rice seedlings under water stress. Rice seedlings were either untreated or subjected to 3 or 24 h of water stress, respectively. Roots were stained with Schiff’s reagent (c) and Evans blue (d), and then immediately photographed under a Leica S6E stereomicroscope (Leica, Solms, Germany). Red/purple indicates the presence of lipid peroxidation detected with Schiff’s reagent. Bar=1 mm. Endogenous NO concentrations and histochemical detection of oxidative damage in the root are given. In Fig. 2b, the red dotted oval represents the high endogenous NO production in the NH_4_^+^- and NO_3_^-^-supplied rice, respectively. Values represent means± standard error (SE) (n=10). CK indicates control treatment, i.e., plants receiving sufficient water.

Histochemical visualization by Schiff’s reagent and Evans blue staining showed that water stress caused severe oxidative damage to the plasma membrane and cell death in the roots of the plants receiving NO_3_^-^, whereas the damage was far less pronounced in the seedlings given NH_4_^+^ (Fig. 1c, d). The following analysis of the MDA and carbonyl concentrations also confirmed that water stress induced more severe lipid peroxidation in the roots of NO_3_^-^-treated than in the roots of NH_4_^+^-treated seedlings.

### Effects of the NO donor on root NO production and oxidative damage

To determine the roles of NO in water stress tolerance, the NO donor SNP was used to simulate NO production. Pre-experimentation with various SNP concentrations (0-100 μM) was performed to quantify the efficacy of SNP against root oxidative damage. As shown in Supplementary Fig. S2, root oxidative damage induced by water stress was significantly alleviated by ≤20 μM SNP. However, the remedial effect of SNP on root oxidative damage was reversed at higher application doses (≥40 μM), suggesting that high SNP or NO contents are toxic to root growth. Therefore, 20 μM SNP was used in the NO donor experiments conducted in the present study. After3 h of water stress, SNP application significantly increased root NO fluorescence intensity for both N treatments. At 3 h, the NO production levels were ~2.05× and 3.85 × higher in the SNP-treated roots of the seedlings receiving NH_4_^+^ and NO_3_^-^, respectively, than in the roots of CK plants (Fig. 2a, b). However, this phenomenon was not observed after 24 h of water stress.

**Fig. 2.**
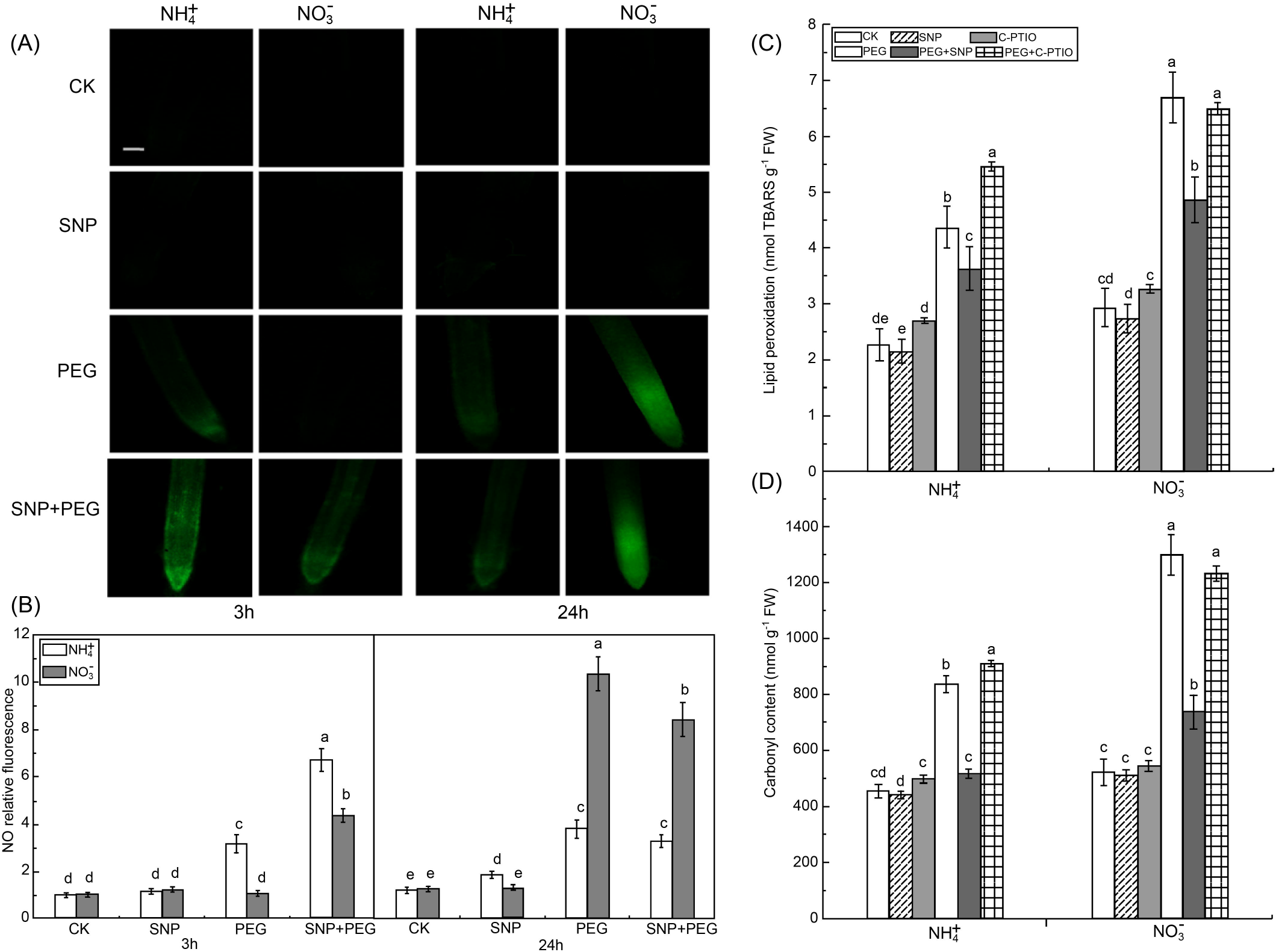
Responses of endogenous nitric oxide (NO) concentrations to water stress (a, b) and NO donor and NO scavenger (c, d) in root apices. (a) Photographs of NO production after sodium nitroprusside (SNP) application. Bar=300 μm. (b) NO production expressed as relative fluorescence. Rice seedlings were either untreated or treated with SNP under water stress. After 3 h and 24 h of treatment, root tips were loaded with 10 μM 4-amino-5-methylamino-2’,7’-difluorofluorescein diacetate (DAF-FM DA) and NO fluorescence was imaged after 20 min using a fluorescence microscope. Endogenous NO concentrations in root are displayed. Values represent means± standard error (SE) (n=10). Different letters indicate significant differences at P<0.05. CK, control treatment, i.e., plants receiving sufficient water.

After 3 h of water stress, ROS (O_2_^-^, -H_2_O_2_, and OH^-^) levels were increased in the roots of both the NH_4_- and NO_3_^-^-treated seedlings in relation to that of CK seedlings. Under water stress, the O_2_^-^, H_2_O2, and OH^-^ in the roots given NH_4_^+^ and NO_3_^-^increased by 78.1% and 107.3%, 28.3% and 47.8%, and 10.6% and 48.4%, respectively (Fig. 3a-c). After 3 h of water stress, root MDA and carbonyl were ~1.28× and 1.4× higher in the plants receiving NO_3_^-^ than in CK plants, respectively. In turn, MDA and carbonyl levels were significantly higher in the CK than in the plants given NH_4_^+^ (Fig. 4a, b). Water stress induced higher root ONOO^-^ in the NH_4+_^-^-treated plants than in the NO_3_^-^-treated seedlings (Fig. 4d), and exogenous NO significantly reduced water stress-induced ROS (O_2_^-^ and H_2_O2) accumulation and oxidative damage (as reflected by MDA and carbonyl) in both N treatments (Figs. 3, 4).

**Fig. 3.**
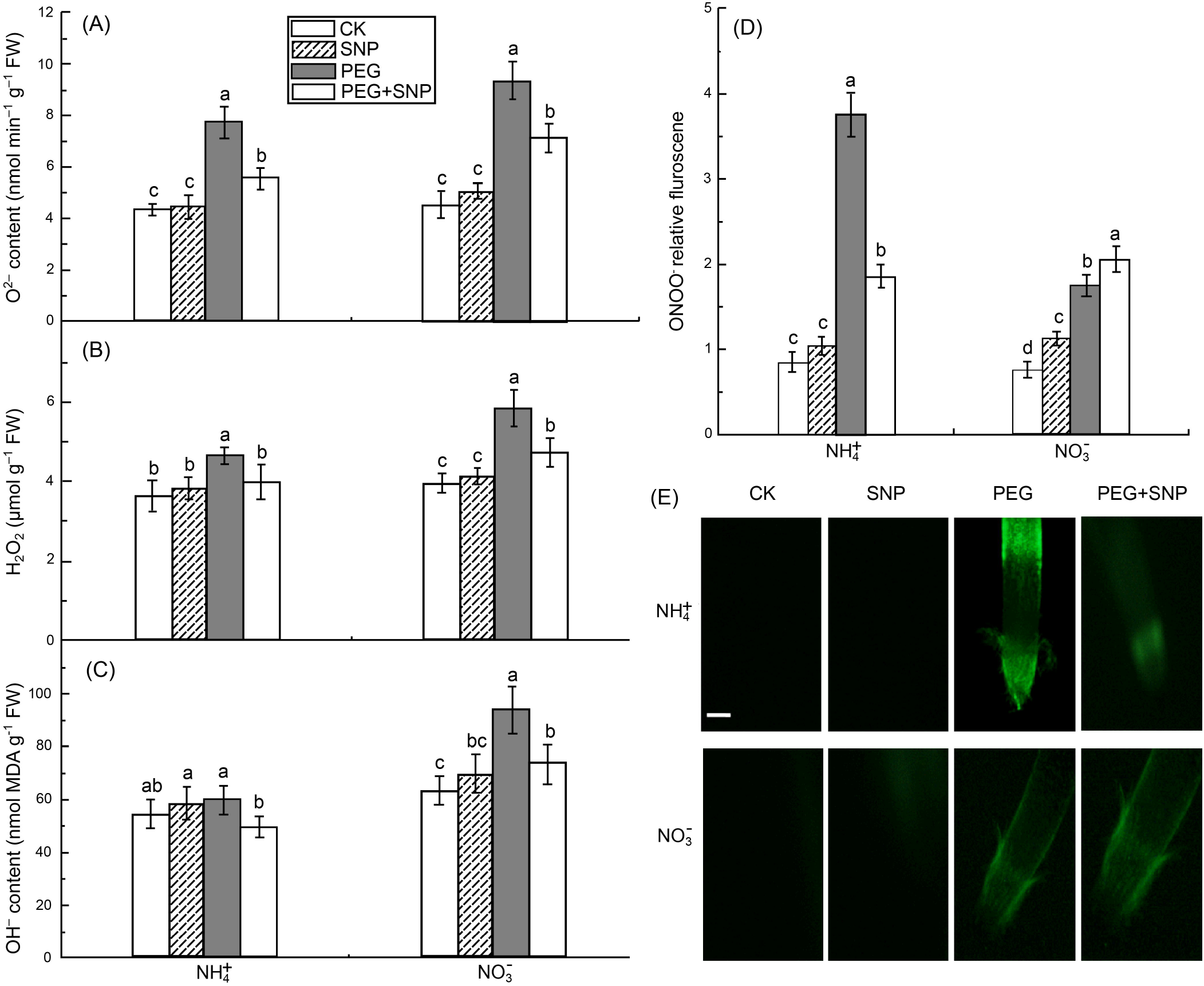
Reactive oxygen species (ROS) and peroxynitrite (ONOO^-^) accumulation in root apices of rice seedlings treated with NO donor (sodium nitroprusside) and either receiving sufficient water (CK treatment) or subjected to water stress using polyethylene glycol(PEG). After 3 h, O^2-^ (a), H_2_O_2_ (b), and OH^-^ (c) levels in rice seedlings roots were measured by spectrophotometry. The accumulation of ONOO^-^was detected with 10 μM aminophenyl fluorescein. Fluorescence images and relative fluorescence intensity were analyzed as described in Fig. 2 for NO determination.

**Fig. 4.**
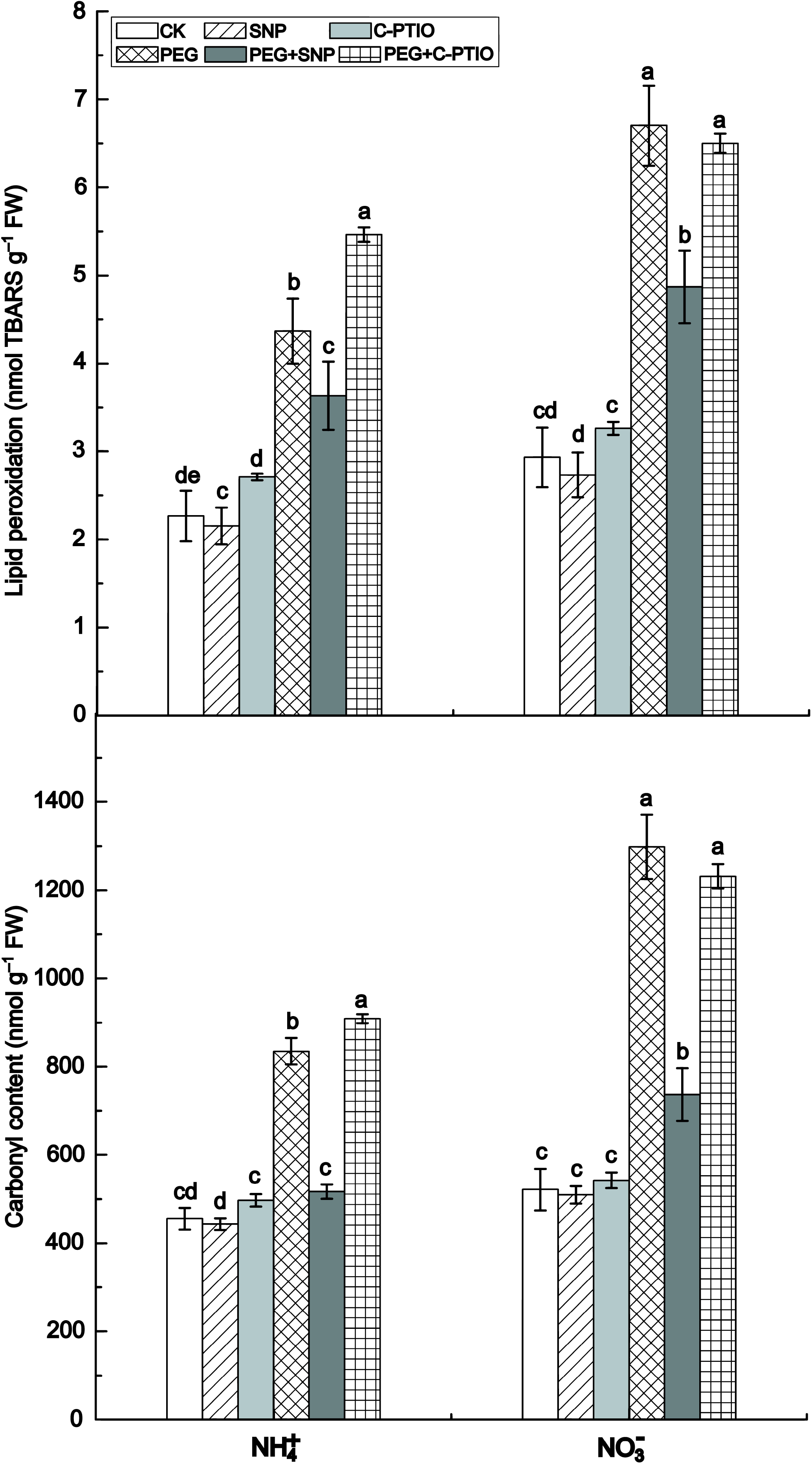
Responses of oxidative damage in root apices of rice seedlings to NO donor (sodium nitroprusside) and NO scavenger (2-(4-carboxyphenyl)-4,4,5,5-tetramethylimidazoline-1-oxyl-3-oxide; c-PTIO) in plants either receiving sufficient water (CK treatment) or subjected to water stress using polyethylene glycol (PEG). In the c-PTIO and PEG + c-PTIO treatments, the rice seedlings were pretreated with NO scavenger (c-PTIO) for 3 h followed by sufficient water or water stress. After 3 h, the malondialdehyde (MDA) content was determined. The MDA in rice roots represents lipid peroxidation (a) and carbonyl concentration (b). Values represent means± standard error (SE) (n=6). Different letters indicate significant differences at *P*<0.05 level.

To determine whether the alleviation of water stress-induced oxidative damage by SNP was related to NO production, the NO scavenger c-PTIO was applied to the plants. After pretreatment with 100 μM c-PTIO for 3 h, alleviation of the water stress-induced root oxidative damage by SNP was reversed (Fig. 4). Depletion of endogenous NO by c-PTIO significantly aggravated root oxidative damage in the NH_4_^+^-treated plants but had no significant effect on the NO_3_^-^-treated plants (Fig. 4), in relation to that observed in CK plants. Therefore, the water stress-induced early NO burst observed in the NH_4_^+^_-_-treated plants alleviates root oxidative damage by reducing ROS, such as O2^-^ and H_2_O_2_^-^.

### Source of endogenous NO

Endogenous plant NO production is mostly driven by NR and NOS. Water stress increased NR activity in the NO_3_^-^-treated roots, and this activity was higher at 24 h than it was at 3 h of water stress (Supplementary Fig. S3a). The activity of NOS was also significantly elevated at 3 h of water stress, and significantly higher in the NH_4_^+^-treated than in the NO_3_^-^-treated roots (Supplementary Fig. S3b). In contrast, water stress suppressed NOS activity in the NO_3_^-^-treated roots at 24 h. Tungstate and L-NAME, which inhibit NR and NOS activities, respectively, were used to identify the origin of the early NO burst in the NH_4_^+^-treated roots. Although L-NAME significantly inhibited endogenous NO production in the NH_4_^+^-treated roots under 3 h water stress, it had no significant effect in the NO_3_^-^-treated roots. At 24 h, the tungstate and L-NAME applications suppressed NO production in the NO_3_^-^-treated roots and tungstate had the stronger inhibitory effect. On the other hand, tungstate had no significant effect on NO production in the NH_4_^+^-treated roots (Fig. 5a, b).

**Fig. 5.**
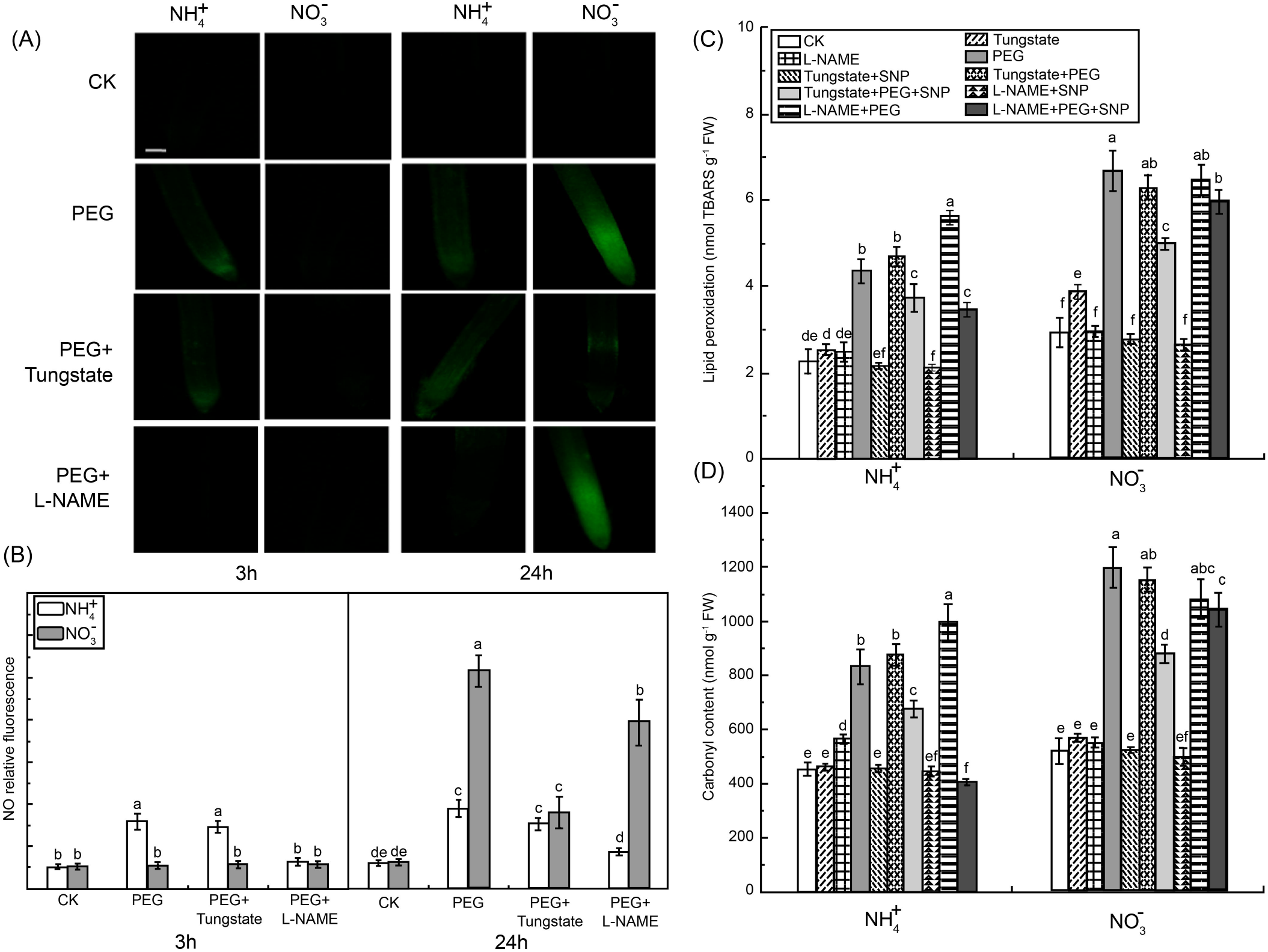
Effects of a nitrate reductase(NR) inhibitor (tungstate) and a nitric oxide (NO) synthase (NOS) inhibitor (Nx-nitro-*L*-arginine methyl ester hydrochloride; L-NAME) on NO content and oxidative damage in root apices of rice seedlings. Rice seedlings were pretreated with NR inhibitor (100 μM tungstate) or NOS inhibitor (100 μM L-NAME) for 3 h, and then subject to water treatment. (a) NO fluorescence. Bar = 300 μm. (b) NO production expressed as relative fluorescence. (c, d) malondialdehyde (MDA) content representing lipid peroxidation (c) and carbonyl concentration (d) in rice seedling roots measured after 3 h of water treatment following tungstate or L-NAME pretreatment. Values represent means± standard error (SE) (n=6). Different letters indicate significant differences at P<0.05 level. CK, control treatment, i.e.,plants receiving sufficient water.

The effect of SNP on the alleviation of water stress-induced root oxidative damage was reversed after pretreatment with 100 μM c-PTIO for 3 h. Application of the NOS inhibitor c-PTIO significantly aggravated water stress-induced oxidative damage in the NH_4_^+^-treated roots, and SNP application reversed the effect of the NOS inhibitor but not that of the NR inhibitor (Fig. 5c, d). For the NO_3_^-^-treated roots, the application of the NR inhibitor or NOS inhibitor had no significant effect on root oxidative damage relative to the PEG (water stress) treatment.

### Activities of antioxidative enzymes and nitrate/nitrite and arginine/citrulline metabolism

Water stress significantly enhanced the activities of root antioxidant enzymes CAT, SOD, APX, and POD by ~107% and 38%, 52% and 36%, 152% and 128%, and 45% and 37% in the NH_4_^+^-treated roots and the NO_3_^-^-treated roots, respectively,compared to the CK roots (Fig. 6). While SNP application further increased CAT, SOD, and APX activities (Fig. 6a-c), these antioxidant enzymes were inhibited by the application of the NO scavenger c-PTIO and by the NOS inhibitor L-NAME in the NH_4_^+^-treated roots under water stress.

**Fig. 6.**
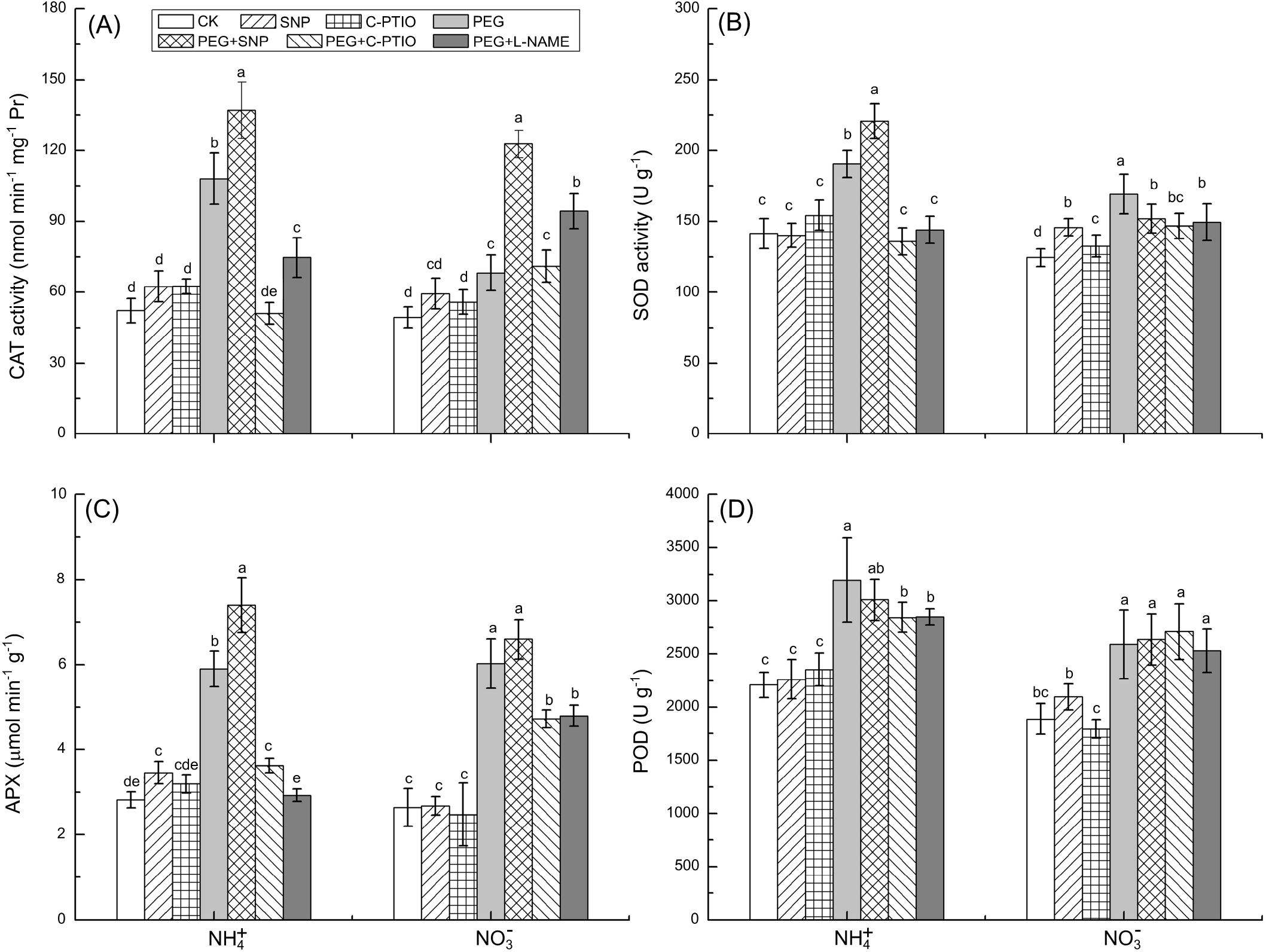
Effects of different treatments on antioxidant enzyme changes in rice seedlings under water stress. Roots were collected to assay catalase (CAT) (a), superoxide dismutase (SOD) (b), ascorbate peroxidase (APX) (c), and peroxidase (POD) (d) after 3 h of treatment with sufficient water (CK treatment) or water stress. For the 2-(4-carboxyphenyl)-4,4,5,5-tetramethylimidazoline-1-oxyl-3-oxide (c-PTIO),polyethylene glycol (PEG) + c-PTIO, and PEG + Nx-nitro-*L*-arginine methyl ester hydrochloride (L-NAME) treatments, the rice seedlings were pretreated with NO scavenger (c-PTIO) or NOS inhibitor (100 μM L-NAME) for 3 h followed by sufficient water or water stress. Values represent means± standard error (SE) (n=6). Different letters indicate significant differences at *P*<0.05 level.

As NR and NOS activities increased in the NO_3_-treated roots, water stress lowered the nitrate level in the NR pathway and the arginine level in the NOS pathway (Supplementary Fig. S4a, b). Similarly, NR inhibitor and NOS inhibitor applications enhanced root nitrate and arginine contents, respectively. In the NH_4_^+^-treated roots, water stress significantly decreased arginine level, indicating that arginine metabolism was relatively high. In this treatment, the NR inhibitor had no significant effect on root arginine content. On the other hand, the NOS inhibitor suppressed arginine metabolism, and thus the NH_4_^+^-treated roots had higher arginine levels than CK roots (Supplementary Fig. S4c). These results indicate that the NO production burst in the NH_4_^+^-treated roots might originate from the NOS pathway.

## Discussion

Ample experimental evidence has demonstrated that NO is involved in plant abiotic stress (Neill *et al.,* 2003; Santisree *et al.,* 2015). However, to our knowledge, no detailed study has been conducted to evaluate the role of NO in drought acclimation in plants supplied with NO_3_^-^ or NH_4_^+^. In the present study, biomass, root N uptake rate, and leaf photosynthesis were reduced relative to the control treatments after 21days of water stress (Supplementary Fig. S1). However, these reductions were less severe for seedlings receiving NH_4_^+^ suggesting that NH_4_^+^ supplementation can enhance drought tolerance in rice seedlings more effectively than NO_3_^-^ supplementation (Guo *et al.,* 2007; Li *et al.,* 2012). Our study also demonstrated that, in the short term (48 h), endogenous NO production in response to water stress is usually time-dependent, varying according to water stress duration. This finding is consistent with those reported for other stressors (Planchet *et al.,* 2014; Sun *et al.,* 2014). Early NO bursts were induced at 3 h of water stress in the roots of NH+-treated seedlings but not in the roots of NO_3_^-^-treated seedlings. Thus, there might be significant differences between NH_4_^+^-/NO_3_^-^-supplied plants in terms of NO signal-mediated drought tolerance. In addition, accumulation of ROS, such as O_2_^-^, OH^-^, and H_2_O2, and root oxidative damage were significantly lower in the NH_4_^+^-treated than in the NO_3_^-^-treated roots at 3 h of water stress. Because ROS accumulation damages cells and their plasma membranes by inducing lipid peroxidation (oxidative stress) (Jiang and Zhang, 2002), the early NO burst in response to water stress observed in NH_4_^+^-supplied seedlings might play a crucial role in their antioxidant defense system and drought tolerance.

The role of the early NO burst in the water stress tolerance of NH_4_^+^-/NO_3_^-^-supplied seedlings was confirmed using NO donors and scavengers. Our study demonstrated that NO donors induced NO in the NO_3_^-^-treated roots at 3 h but not at 24 h of water stress. Plant ROS accumulation and MDA and carbonyl levels under water stress were significantly alleviated after the application of the NO donor in both N treatments. Nevertheless, the levels of these substances were higher in the NO_3_^-^-treated roots than in the NH_4_^+^-treated roots. Therefore, the NO production enhanced at 3 h by the exogenous NO donor can alleviate water stress-induced oxidative damage in the NO_3_^-^-treated roots. On the other hand, elimination of the early NO burst by NO scavengers like c-PTIO significantly aggravated water stress-induced oxidative damage. These results provide direct evidence that the early NO bursts plays a crucial role in drought tolerance in NH_4_^+^-treated roots. Because theNH_4_^+^-supplied roots maintained a higher N uptake rate thanNO_3_^-^-supplied roots under water stress (Supplementary Fig. S1f), we hypothesized that the higher NH_4_^+^ uptake rate is beneficial for the NO early burst due to the NO production involved in root N metabolism (Corpas *et al.,* 2008; del Rio, 2015). This NO burst can also be an active adaptation mechanism of plants to abiotic stress as, in addition to drought stress, it has been reported to occur repeatedly in plants challenged by pathogens (Floryszak-Wieczorek *et al.,* 2007), metal toxicity (Gonzalez *et al,* 2012; Sun *et al.,* 2014), and cold stress (Cantrel *et al.,* 2011).

Our study demonstrated that an early NO burst improves plant drought tolerance by enhancing the antioxidant defense system of the root. Elevated plant antioxidant enzyme activities and gene expression levels in response to water stress have been widely demonstrated (Jiang and Zhang, 2002; Arasimowicz-Jelonek *et al.,* 2009a; Fan and Liu, 2012). In the present study, the tips of the NO_3_^-^-treated roots presented more serious water stress-induced oxidative damage (due to the excessive production ofO_2_^-^, OH^-^, and H_2_O2) than those of the NH_4_^+^-treated roots (Figs. 1-3). In contrast, NH_4_^+^-supplied roots maintained relatively higher antioxidant enzyme (CAT, SOD, and APX) activity levels to catalyze O2^-^ and H_2_O2 decomposition (Fig. 3). It has been demonstrated that there is significant crosstalk between NO and ROS in plants. The antioxidant function of NO was explained by its ability to reduce H_2_O2 and lipid peroxidation, and induce antioxidant gene expression and enzyme activity (Bogeat-Triboulot *et al.,* 2007; Farooq *et al.,* 2009). Our results showed that enhanced NO levels and antioxidant enzymeactivities(CAT and SOD) were significantly and simultaneously increased after NO donor application in NO_3_^-^-treated roots thereby reducing ROS concentration and oxidative damage. The early NO burst observed in NH_4_^+^-treated roots can enhance antioxidant enzyme activity and ROS accumulation (O_2_^.-^, OH_2_^-^, and H_2_O_2_). These results were confirmed by subsequent experimentation in which the application of NO scavenger significantly suppressed SOD and CAT in NH_4_^+^-treated roots. Thus, drought tolerance in the NH_4_^+^-treated roots might be associated with the NO induced up-regulation of antioxidant enzymes and down-regulation of ROS accumulation.

Nitric oxide can also serve as a source of reactive nitrogen species (RNS). Over accumulation of RNS under abiotic stress can cause tyrosine nitration and inactivate proteins like CAT, manganese-dependent (Mn-)SOD, and GR (Clark *et al.,* 2000) as well as the peroxidative activity of cytochrome c (Batthyany *et al.,* 2005). Our results show that NO_3_^-^-supplied plants had more severe oxidative damage and accumulated extremely high NO levels after 24 h of water stress. This latent NO production can be partially alleviated by replenishing the early NO burst at 3 h with SNP (Fig. 1). These results indicate that both ROS and RNS metabolism participate in the water stress response. High NO accumulation in the NO_3_^-^-treated roots likely cause the nitrosative stress at 24 h, which also damaged root redox balance. A similar phenomenon was described in plants subjected to cold (Airaki *et al.,* 2012), salinity (Tanou *et al.,* 2012), and drought (Signorelli *et al.,* 2013) stresses. Because NO competes with oxygen for cytochrome c oxidase binding (Complex IV), it affects both the respiratory chain and oxidative phosphorylation (Millar and Day, 1996; Yamasaki *et al.,* 2001). Thus, under drought stress, the higher NO production in the NO_3_^-^-treated roots than in the NH_4_^+^-treated roots could aggravate respiratory inhibition and induce greater oxidative damage.

Our investigation suggests that the early NO burst in NH_4_^+^-treated roots is mainly mediated NOS at the early stages of water stress. Nitrate reductase-mediated NO generation is known to occur under water deficit (Arasimowicz-Jelonek *et al.,* 2009b; Yu *et al.,* 2014). Drought-induced NO generation by NOS-like enzymes in plants has also been demonstrated but this NO production pathway varies significantly with species, tissue type, and plant growth conditions (Corpas *et al.,* 2009; Liao *et al.,* 2012; Shi *et al.,* 2014). For the NH_4_^+^-treated roots, both NOS activity and NO production increased simultaneously at 3h of water stress, whereas the application of the NOS inhibitor completely repressed NO synthesis at this time point. The NOS inhibitor also aggravated water stress-induced membrane lipid peroxidation and oxidative protein damage, indicating that some NOS associated proteins may play an important role in NO-mediated drought protective responses (Guo *et al.,* 2003; Zhao *et al.,* 2007). In contrast, the NR inhibitor did not significantly affect NO production or membrane lipid peroxidation. The aggravation of lipid peroxidation by L-NAME may have been the result of the alteration of the NOS-mediated early NO burst. In NO_3_^-^-treated roots, water stress enhanced NR activity significantly more than NOS activity at 24 h. However, separate NR inhibitor and NOS inhibitor applications only partially suppressed NO production. The NO produced by the NR pathway might therefore play an important role in later NO production (24 h), consistent with previous reports (Arasimowicz-Jelonek *et al.,* 2009a, b). Although several studies support the arginine-dependent NO production model in higher plants, the genes encoding NOS in such plants have not yet been identified (Zemojtel *et al.,* 2006). For this reason, the nitrate/nitrite and arginine/citrulline levels in the NR and NOS pathway, respectively, were determined. It was found that water stress significantly increased NOS activity and accelerated the conversion of arginine to citrulline. However, the arginine content was significantly enhanced in the NH_4_^+^-treated roots after the NOS inhibitor application, in relation to the CK roots. These results provide additional evidence that the early NO burst in NH_4_^+^-treated roots is mainly mediated by NOS (Fig. 7).

**Fig. 7.**
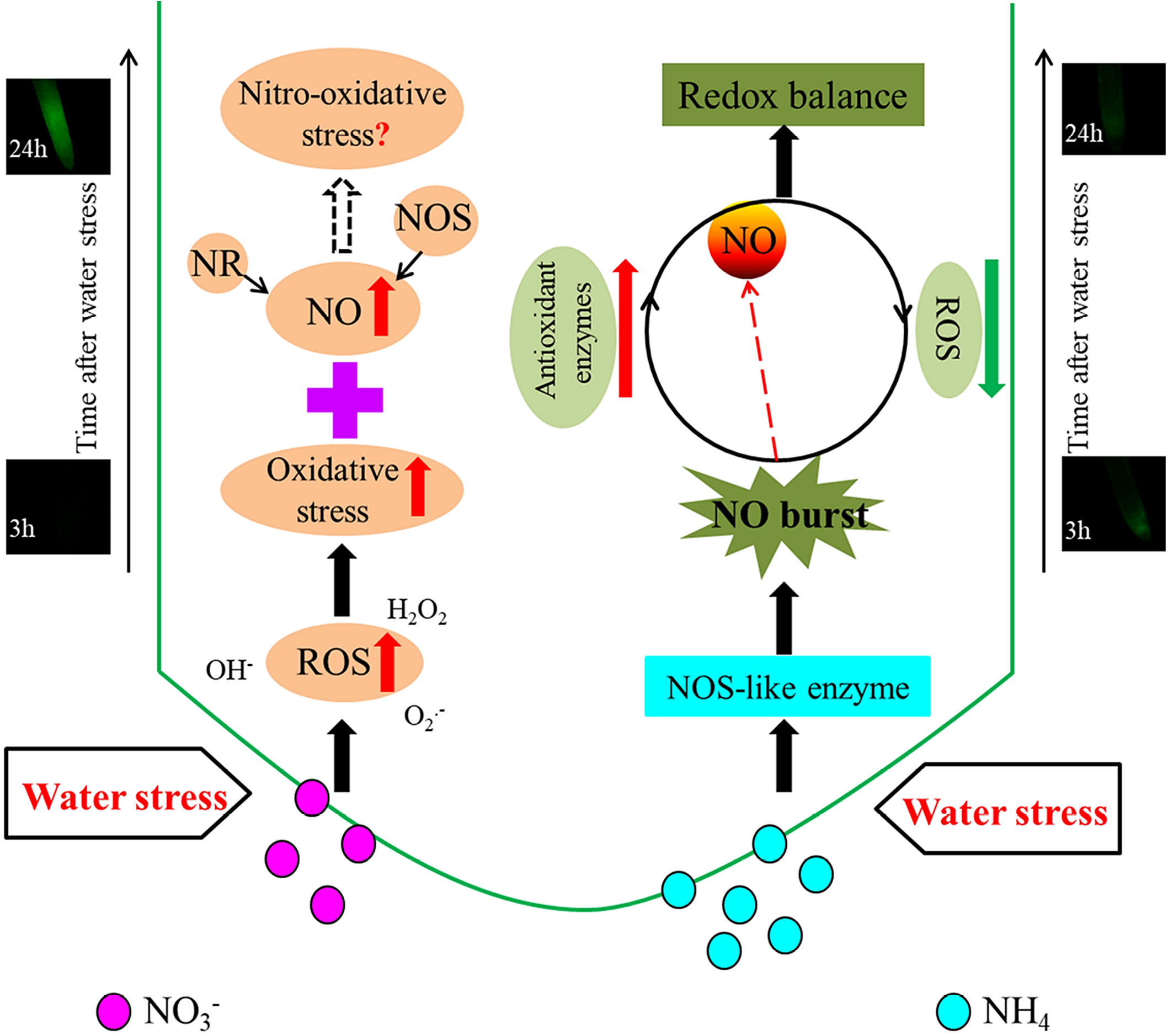
Schematic illustration of a proposed model for the different responses of early NO production and its effects on the defense response of rice to water stress. In the roots of NH_4_^+^-supplied rice, the nitric oxide synthase (NOS)-mediated early nitric oxide (NO) burst (3h) significantly enhanced plant antioxidant defense by reducing reactive oxygen species (ROS) accumulation and enhancing antioxidant enzyme activity; the relative lower NO production after 24 h of water stress in comparison to NO_3_^-^-supplied rice, also helped maintaining the redox balance in root cells, thus enhancing their drought tolerance. In the roots of NO_3_^-^-supplied rice, ROS accumulation and oxidative damage induced by 3h of water stress were significantly higher than that in NH_4_^+^-supplied rice. High NO accumulation in the NO_3_^-^-treated roots likely caused the nitrosative stress at 24 h of water stress. A combined effect of oxidative and nitrification stresses might have led to the weak resistance to water stress in NO_3_^-^-supplied rice. NR, nitrate reductase. Red arrows represent increase, green arrows represent decrease. Black solid arrows represent defined pathways, dotted arrows represent undefined pathway.

Our study is the first to demonstrate that the early NO burst in NH_4_^+^-treated rice roots significantly enhanced plant antioxidant defense by reducing ROS accumulation and enhancing the activities of antioxidant enzymes, thereby increasing plants’ acclimation to water stress. The early NO burst which occurs in response to water stress may be triggered by NOS-like enzymes in root. Our results provide new insight into how NO-signaling molecules modulate drought tolerance in NH_4_^+^-supplied rice plants. However, the signaling crosstalk between ROS and RNS in response to water stress merits further investigation and may help elucidate the role of the NO-signaling process in enhancing drought tolerance in NH_4_^+^-supplied rice.

## Supplementary data

**Fig. S1**. Responses to water stress in NH_4_^+^- and NO_3_^-^-supplied rice

**Fig. S2**. Effects of exogenous nitric oxide donor on root oxidative damage under water stress

**Fig. S3**. Effect of water stress on nitrate reductase and nitric oxide synthase in rice roots

**Fig. S4**. Levels of nitric oxide-related compounds under sufficient water or water stress

## Method S1

Determination of leaf photosynthesis, root N uptake rate, root nitrate and nitrite content in rice seedlings after 21days under control or water stress treatments

## Acknowledgements

This work was supported by the National Key Research and Development Program of China (No. 2017YFD0300100, 2016YFD0101801); the Natural Science Foundation of Zhejiang Province (No. LY18C130005). We would like to thank Editage [www.editage.cn] for English language editing.

---

**Fig. S1.** (a), (b), and (c) response of NH_4_^+^- and NO_3_^-^-supplied rice agronomic characteristics and biomass to water stress induced by 10% PEG after 21 days of treatment. (d) Effects of water stress on leaf photosynthesis in NH_4_^+^- and NO_3_^-^-supplied rice after 21days of treatment. (e) Effects of water stress on root activity in NH_4_^+^- and NO_3_^-^-supplied rice after 21days of treatment. (f) Effects of water stress on root ^15^N-labeled uptake rate in NH_4_^+^- and NO_3_^-^-supplied rice after 21days of treatment. Rice leaf photosynthesis, root activity, and ^15^N uptake rate were determined according to Method S1. Values represent means±SE (n=6). Different letters indicate significant differences at P<0.05 level. CK, control treatment, i.e., plants receiving sufficient water.

**Fig. S2.** Effect of exogenous NO donor (SNP) on root oxidative damage under water stress. Rice roots were exposed to mixed N (NH_4_^+^ + NO_3_^-^) nutrient solution containing 0 μM, 5 μM, 10 μM, 20 μM, 40 μM, 80 μM, or 100 μM SNP either with or without 10% PEG for 48 h. MDA levels representing lipid peroxidation (a) and carbonyl concentration (b) in rice seedling roots were determined. Values represent means±SE (n=6). Different letters indicate significant differences at P<0.05 level. CK, control treatment, i.e., plants receiving sufficient water.

**Fig. S3.** Effect of water stress on NR (a) and NOS (b) in root apices of rice seedlings. Roots were collected for the NR and NOS assays after 3 h and 24 h of water stress, respectively. Values represent means±SE (n=6). Different letters indicate significant differences at P<0.05 level. CK, control treatment, i.e., plants receiving sufficient water.

**Fig. S4.** Related compounds in NR-mediated and NOS-mediated NO pathways in root apices of rice seedlings treated with NR inhibitor (tungstate) and NOS inhibitor (L-NAME) under sufficient water or water stress treatment. (a) Levels of nitrate and nitrite in NO_3_^-^-treated roots. (b) Levels of arginine and citrulline in NO_3_^-^-treated roots.(c) Levels of arginine and citrulline in NH_4_^+^-treated roots. For the PEG + Tungstate and PEG + L-NAME treatments, the rice seedlings were pretreated with NR inhibitor (100 μM tungstate) or NOS inhibitor (100 μM L-NAME) for 3 h, followed by sufficient water (CK treatment) or water stress treatment. Values represent means±SE (n=6). Different letters indicate significant differences at P<0.05 level.

